# Genome Wide Association Study of Resistance to PstS2 and Warrior Races of Stripe (Yellow) Rust in Bread Wheat Landraces

**DOI:** 10.1101/2020.02.09.940775

**Authors:** Muhammad Massub Tehseen, Fatma Aykut Tonk, Muzaffer Tosun, Ahmed Amri, Carolina P. Sansaloni, Ezgi Kurtulus, Mariana Yazbek, Khaled Al-Sham’aa, Izzet Ozseven, Luqman Bin Safdar, Ali Shehadeh, Kumarse Nazari

## Abstract

Stripe rust, caused by *Puccinia striiformis* Westend. f. sp. *tritici* is a major threat to wheat production worldwide. The breakdown in resistance of certain major genes and new emerging aggressive races of stripe rusts are causing serious concerns in all main wheat growing areas of the world. To search for new sources of resistance genes and associated QTL for effective utilization in future breeding programs an association mapping panel comprising of 600 bread wheat landraces collected from eight different countries conserved at ICARDA gene bank were evaluated for seedling and adult plant resistance against *PstS2* and *Warrior* races of stripe rust at the Regional Cereal Rust Research Center (RCRRC), Izmir, Turkey during 2016, 2018 and 2019. A set of 25,169 informative SNP markers covering the whole genome were used to examine the population structure, linkage disequilibrium and marker-trait associations in the association mapping panel. The genome-wide association study (GWAS) was carried out using a Mixed Linear Model (MLM). We identified 47 SNP markers at 19 genomic regions with significant SNP-trait associations for both seedling and adult plant stage resistance, the threshold of significance for all SNP-trait associations was determined by the false discovery rate (q) ≤ 0.05. Three genomic regions (*QYr.1D_APR, QYr.3A_seedling* and *QYr.7D_seedling*) identified in this study are far away from any previously reported *Yr* gene or QTL hence, tagging novel genomic regions. The *In-silico* analysis of the novel QTL regions identified candidate resistance genes encoding proteins putative to plants disease resistance and defense mechanism.

## INTRODUCTION

Wheat stripe rust, caused by *Puccinia striiformis* Westend. f. sp. *tritici* (*Pst*), is one of the most devastating diseases of wheat worldwide. Historically, the stripe rust is considered a disease of cool and wet climates but has recently shifted towards warmer climatic regions (Muleta et al., 2017). Stripe rust epidemic has reported causing grain yield loss up to 40% to 100% under severe infections (Manickavelu et al., 2016; Mumtaz et al., 2009). The emergence of new aggressive and high-temperature tolerant pathotypes of stripe rust has changed the epidemiology of the pathogen and now the disease is developing in areas once considered unfavorable for stripe rust infestation (Hovmoller et al., 2011; Sorensen et al., 2014). These new pathotypes of *Pst* are currently widespread from Asia to Europe, Africa and Australia threating wheat yields at a global level.

The continuous emergence of new pathotypes of *Pst* creates a need to develop new cultivars and strategies to resist the epidemic through time. The most economical and effective way of controlling stripe rust outbreaks is the use of genetic resistance equipped with both minor and major genes (Chen 2013). Two types of genetic resistance based on major and minor genes are deployed in the breeding programs all over the world (Chen 2005). Resistance carried out by major genes is often referred to as race-specific, seedling and/or all stage resistance, it is based on “R” genes, and is effective at all stages of plant life (Burdon et al., 2014). The commercial cultivars with resistance controlled by R genes are short-lived as new virulent pathotypes continuously emerge in the pathogen population eventually overcoming the R genes. Many devastated epidemics around the world are the result of the transient nature of the R gene-mediated resistance in the commercial cultivars (Ellis et al., 2014; Hulbert & Pumphrey 2014; Steele et al., 2001). On the other hand, minor gene resistance is usually expressed at the adult plant stage and usually referred to as Adult Plant Resistance (APR). In many cases, minor gene or APR resistance is considered more effective and durable. Due to the rapid evolution of *Pst* pathogen, many virulent pathotypes have emerged therefore, the best strategy is to pyramid both major and minor genes for more durable rust resistance (Singh et al. 2000; Singh et al., 2005).

*PstS2* and *Warrior* (*PstS7*) races are the two most widely spread pathotypes of *Pst* covering the geographical regions from Asia all the way to Northern Europe (Tadesse et al., 2014; Hovmoller et al., 2016). *Warrior* was first discovered in the UK in 2011, and currently is the most prevalent race of *Pst* in Europe (Ali et al., 2017) and it was detected in Turkey in 2014 (Mert et al., 2015). The breakdown of resistance gene *Yr27* in many countries in Central West Asia and North Africa (CWANA) region caused yield losses of up to 80% (Solh et al., 2012). The development of genetic material resistant to these two prevalent races of *Pst* is very crucial for the region. Deployment of the combination of major and minor genes is the long-term goal of most of the breeding programs around the globe. This strategy provides resistance to broader spectrum of races thus providing wider durability against multiple virulent pathotypes (Chen 2013; Singh et al., 2000). Recent advances in genomics and statistical approaches that provide effective marker tagging systems have led to facilitate the strategy to pyramid major-minor genes with higher efficiency using advanced means of genome manipulation (Muleta et al.,. 2017; Bentley et al.,. 2014).

Genome-Wide Association Studies (GWAS) takes advantage of historical recombination since population diversification by utilizing Linkage Disequilibrium (LD) within the gene pool of a species to identify phenotype-genotype correlations and is widely used to detect quantitative trait loci (QTL) in plants (Muleta et al.,. 2017; Manickavelu et al., 2016 Bulli et al., 2016; Maccaferri et al., 2015; Kertho et al., 2015). Furthermore, GWAS detects QTL with higher resolution from germplasm of diverse background therefore, effectively eliminating the time and cost spent in the development of bi-parental mapping populations. The primary gene pool of wheat preserved at the gene banks includes landraces, traditional cultivars and breeding lines offering a diverse range of sources of resistance to biotic and abiotic stresses including disease resistance (Endresen et al.,. 2011). The co-evolution of landraces along with biotic and abiotic stresses has made them an important choice in the selection of breeding/pre-breeding materials for crop improvement (Zeven 1998). Similarly, landraces have co-evolved with rust pathogens which might have resulted in the accumulation of diverse resistance loci. These landraces may possess useful untapped genetic resistance loci since they have not been widely utilized in the breeding programs. The use of germplasm preserved at gene banks for disease resistance in wheat has been previously reported in numerous studies (Muletta et al., 2017; Bulli et al., 2016; Manickavelu et al., 2016; Maccaferri et al., 2015; Kertho et al., 2015; Naruoka et al., 2015; Gurung et al., 2014).

In this study, we examined the hypothesis that if bread wheat (*Triticum aestivum* L.) landraces preserved at The International Center for Agriculture Research in the Dry Areas (ICARDA) gene bank can provide a useful source of resistance against critically important *Pst* race in CWANA. The present study has the following three objectives: (1) to evaluate the diversity for seedling and adult-plant resistance to *Pst* pathotypes *PstS2* and *PstS7* (*Warrior*) in bread wheat landraces, (2) to perform GWAS analysis to identify single-nucleotide polymorphism (SNP) loci associated with resistance to these two Pst pathotypes, (3) to compare and evaluate the *Pst* resistance loci identified in this study with previously identified *Yr* genes and QTLs, (4) to perform *in-silico* analysis to search for the presence of candidate genes with putative disease resistance proteins in association with the loci identified in this study and previously identified *Yr* genes and QTL.

## MATERIALS AND METHODS

### Plant Material

A panel of 600 bread wheat landraces preserved at the ICARDAs gene bank was selected for this study. This panel was a collection of landrace accessions from Syria (376 accessions), Turkey (157), Iran (47), Greece (7), Iraq (7), Spain (3), Jordan (2) and Palestine (1). The association mapping panel was evaluated for seedling and APR against *PstS2* and *Warrior* pathotypes of *Pst*. The complete list of landrace accessions and the country of origin is presented in **Supplemental Table S1.**

**Table 1:** Basic statistics of seedling and adult plant response of bread wheat landraces against PstS2 and Warrior pathotypes of stripe rust, estimates of variance components and heritability

| Parameter | Seedling<br>PstS2 | Seedling<br>Warrior | IZM16 | IZM18 | IZM19 | Combined |
| --- | --- | --- | --- | --- | --- | --- |
| Minimum | 1.0 | 1.0 | 2.0 | 1.0 | 1.0 | 1.0 |
| Mean | 6.8 | 7.2 | 29.7 | 31.8 | 26.0 | 29.1 |
| Maximum | 9.0 | 9.0 | 90.0 | 100.0 | 100.0 | 100.0 |
| $\sigma^2_G$ | | | 591.5** | 1048.1** | 788.1** | 178.83** |
| $\sigma^2_E$ | | | 32 | 20.1 | 22.8 | 17 |
| $\sigma^2_P$ | | | 623.5 | 1068.2 | 810.9 | 195.83 |
| Heritability |  |  | 0.95 | 0.98 | 0.97 | 0.91 |

\*Significance at the 5% probability level
\*\* Significance at the 1% probability level
$\sigma^2_G$ = estimates of genotypic variance
$\sigma^2_E$ = estimates of environmental variance
$\sigma^2_P$ = estimates of phenotypic variance

### Seedling stage stripe rust assessment

Association mapping panel was assessed for seedling resistance against the two Pst pathotypes *PstS2* and *Warrior* which were previously collected from Turkey and characterized at the Cereal Rust bio-safety laboratory (BSL) of RCRRC. The virulence/avirulence formulae of the two races along with the differential set used in the study are given in **Supplemental Table S2 and S3**. Eight to 10 seeds of each accession were planted to be tested in BSL against each race of stripe rust races (*PstS2* and *Warrior*). Seeds were planted in 7 cm x 7 cm x 7 cm plastic pots. The mixture in the pots consisted of soil, compost, and sand in a ratio of 1: 1: 1. Seedlings were grown at 17-20 ° C for 10-12 days in a spore proof growth chamber. Inoculation was carried out when the first leaf was fully expanded and the second leaf was half emerged. *PstS2* and *Warrior* urediospores of stripe rust suspended in the light mineral oil (Soltrol 170) were sprayed onto the seedlings with an atomizer. Inoculated seedlings were allowed to dry for few minutes followed by fine misting by distilled water and placing in wet plastic cages containing water at the bottom. Seedlings were then incubated in a dark room for 24 hours at 8-10^◦^C with relative humidity ranging close to 100%. Seedlings were then transferred to a growth chamber with a temperature regime of 15°C for 16 hours’ light at 250 µmol and 8 hours of dark at 10°. Disease assessment was performed 15 days after inoculation using the 0-9 scale (McNeal et al., 1972) for seedling infection types. The range of seedling infections between 0 and 6 were considered low infection types. Seedling infection 7, 8, and 9 were considered as high infection types. The low and high infection types were converted to 0 and 1, respectively when data were used for AM analysis.

**Table 2:** Genomic regions significantly associated with seedling resistance to stripe rust in 600 bread wheat landraces collection from ICARDA gene bank

| SNP<sup>a</sup> | Trait | Chr | Position (bp)<sup>b</sup> | -log10p | R<sup>2</sup> | Previously mapped *Yr* gene/QTL<sup>d</sup> |
| --- | --- | --- | --- | --- | --- | --- |
| 1110537 | Warrior | 1B | 1.35E+08 | 3.52* | 0.047 | *Yr15, YrCH52, Yr64, YrAlp* |
| 100467667 | PstS2 | 2A | 1.44E+08 | 3.64* | 0.074 | *Yrxy2, QTLs QYr.ucw-2A.3, QYr.sun-2AS_Kukri* |
| 100462582 | Warrior | 2A | 6.49E+08 | 3.47* | 0.047 | *QYr.inra_2AL.2_CampRemy* |
| 2272062 | Warrior | 2D | 5.17E+08 | 3.36* | 0.046 | *QYr.tam_2D_Quaiu, QYr.jic-2D_Briagdier* |
| **2307897** | Warrior | 3A | 7.32E+08 | 3.71* | 0.049 |  |
| **5411780** | Warrior | 3A | 7.49E+08 | 3.36* | 0.046 |  |
| 1102819 | Warrior | 3A | 6.47E+08 | 3.64* | 0.048 | *QYrdr.wgp-3AL (IWA6834), QYr.wsu-3A (IWA3401)* |
| 1714660 | Warrior | 3B | 23162218 | 3.59* | 0.048 | *wPt-800213-IWA6510* |
| 1140683 | PstS2 | 3D | 3.31E+08 | 3.37* | 0.072 | *YR45* |
| 1228502 | Warrior | 3D | 5.56E+08 | 3.62* | 0.048 | *IWA3012_seedling* |
| 3957546 | Warrior | 3D | 3.69E+08 | 3.37* | 0.046 | *YR45* |
| 1013303 | PstS2 | 4B | 4.14E+08 | 3.51* | 0.073 | *Yr50, Yr62 QYr.sun-4B_Janz* |
| 1002449 | Warrior | 4B | 6.23E+08 | 3.91** | 0.05 | *QYr.wpg-4B.2 (IWA3994)* |
| 1090007 | Warrior | 5B | 9075119 | 4.09** | 0.052 | *Yr47* |
| 2252899 | Warrior | 5D | 4.07E+08 | 4.13** | 0.052 | *Yr.caas-5DS_Jingshuang 16* |
| 3935577 | PstS2 | 6A | 5.8E+08 | 4.1** | 0.077 | *QYr.cim-6AL_Francolin* |
| 2262271 | PstS2 | 6A | 5.4E+08 | 3.33* | 0.071 | *QYr.caas-6AL (IWB39473)_Zhong 892* |
| 991646 | PstS2 | 6B | 6.57E+08 | 3.72* | 0.074 | *QYr.ucw-6B (IWA7257)* |
| 4990334 | Warrior | 6B | 23343483 | 3.39* | 0.046 | *Yr35, QYr.ufs-6B_Kariega* |
| 999763 | Warrior | 6D | 3.86E+08 | 4.82** | 0.053 | *QYr.ufs-6D_Cappelle-Desprez* |
| 100553682 | PstS2 | 7A | 1.31E+08 | 4.09** | 0.077 | *Yr61, QYr.caas-7A_Jingshuan16* |
| 1695008 | Warrior | 7A | 2.44E+08 | 3.76* | 0.049 | *Yr61, QYr.sun-7A_CPI133872* |
| **2245601** | PstS2 | 7D | 6.02E+08 | 3.56* | 0.073 |  |

<sup>a</sup> SNP index from the GBS data.
<sup>b</sup> Physical positions based on the IWGSC RefSeq. 1.0.
<sup>c</sup> Significance. Experiment-wise (Benjamini-Hochberg FDR adjusted): \* $FDR(q) \leq 0.05$ , \*\* $FDR(q) \leq 0.01$ .
<sup>d</sup> References are given in the text.

### Field Stripe rust assessment

The field experiments were carried out at RCRRC during cropping season 2016 (IZM16), 2018 (IZM18) and 2019 (IZM19). The accessions were planted in 1-meter rows with 30cm spacing between the rows. To ensure sufficient inoculum production for disease infection, a mixture of universal cultivars Morocco and Avocet’s’ along with local wheat susceptible lines were planted as spreader after every 20 rows as well as spreader rows bordering the nurseries. Experiments were managed as per the standard local agronomic practices during the crop season.

Adult-plant resistance of the 600 accessions was evaluated under field conditions against the *PstS2* and *Warrior* pathotypes over three years. *PstS2* and *Warrior* pathotypes collected from previous years and preserved at RCRRC were multiplied at the BSL using susceptible cultivar Avocet’S’. Collected fresh uredeospores were used in field inoculation. The association mapping panel along with spreader rows bordering the experiment were artificially sprayed with the mixture of two races in talcum powder using backpack sprayer at seedling, tillering and booting stages. The field was irrigated through a mist irrigation system.

Field scoring started when the disease severity reached to 100% on susceptible check Morocco and Avocet ‘S’. Adult-plant responses were recorded three times with 10 days’ intervals for the major infection types R, MR, MS, and S (Rolefs et al., 1992) and the disease severities (0-100%) following the Modified Cobb’s Scale (Peterson et al., 1948). All three recordings were averaged and the Coefficient of Infection (CIs) were calculated for infection types and disease severities following Saari & Wilcoxson (1974).

### DNA extraction

Genomic DNA was extracted from fresh leaves collected from three individual 10-days old seedling using a modified CTAB (cetyltrimethylammonium bromide) method (Hoisington et al., 1994). DNA quality and concentration were determined by electrophoresis in 1% agarose gel. Young leaves were collected in labeled Eppendorf tubes and stored in liquid nitrogen at -80°C for DNA extraction. Leaf samples were ground using tissue lyser (Tissue Lyser II from QIAGEN) till fine powder was obtained. 0.1g of the powdered leaf samples were used for DNA extraction using CTAB (Cetyl Trimethylammonium Bromide) method (Doyle, 1990). The extracted DNA was dissolved in 100µl tris-EDTA (TE) buffer. The samples were analyzed on 1% agarose gel for purity and quantified with a spectrophotometer (NanoDrop ND1000). The DNA samples were then kept at -80°C.

### Genotyping

A high-throughput genotyping by sequencing (GBS) method called DArTseq^TM^ technology (Sansaloni et al., 2011) was applied to all samples at the Genetic Analysis Service for Agriculture (SAGA) at the International Maize and Wheat Improvement Center (CIMMYT) in Mexico and supported by the CGIAR Research Program (Sansaloni et al., 2011).

### Population Structure, Linkage Disequilibrium (LD) and Kinship Analysis

To analyze the genetic variation within the population, 1072 unlinked SNP markers were used using STRUCTURE software (v.2.3.4), which implements a model-based Bayesian cluster analysis (Pritchard et al., 2000). The STRUCTURE software divides the populations into clusters to form Q-matrix. The program was operated for ten independent runs for robust evaluation with a putative number of sub-populations ranging from k= 1-10 assessed with a burn-in period of 50,000 steps followed by 50,000 recorded Markov-Chain iterations. The best K value representing the optimum number of clusters in the populations was estimated as Delta K (ΔK) based on the rate of change in log probability of data between successive values using Structure Harvester as described by Evanno et al., (2005). Structure analysis was performed multiple times with altering parameters and iterations to reduce clustering error. Population structure was also analyzed by Principal Component Analysis (PCA). PCA was calculated using R software package PCAdapt which implements method as described by Luu et al., (2017). LD among the markers was estimated for the Association Mapping (AM) panel in Tassel (v.5.2.24) (Bradbury et al., 2007) using the observed vs. expected allele frequencies. The LD decay was measured as the distance at which the average *r^2^* between pairwise SNPs dropped to half of its maximum value (Huang et al., 2010). Tassel (v.5.2.24) was used to derive the population kinship matrix based on the scaled IBS (identity by state) method using the complete set of markers that passed quality filtering as reported by Gao *et al* et al., 2016.

### Association Mapping Analysis

A set of high-quality SNP markers was used for association analysis. Since both model-based Bayesian and PCA revealed a population structure in the panel therefore marker trait association were carried out based on the Mixed Linear Model (MLM Q+K) which accounts for population structure (Q) and kinship matrix (K) using Genome Association and Prediction Integrated Tool (GAPIT) (Lipka et al., 2012) under open-source R environment. Significant markers were identified based on estimating False Discovery rate (FDR) values for each experiment (Ozkuru et al., 2018; Benjamini & Hochberg, 1995). Markers with a minimum threshold of experiment-wise FDR (q) ≤ 0.05 were considered significant. A previously developed integrated map (Bulli et al. 2016) was used to determine the relationships of the SNPs identified in this study with previously reported Yr genes and QTL. Names assigned to the novel QTL identified in this study start with the prefix “Q” for QTL, followed by “Yr” for yellow rust, chromosome name, and “seedling” for seedling trait, “APR” for adult plant resistance.

### Putative candidate gene identification

Candidate genes with their putative proteins/enzymes associated with significant loci from GWAS were predicted on the basis of LD using the International Wheat Genome Sequencing Consortium (IWGSC) *RefSeq v1.0* annotations (Appels et al., 2018) available at https://wheat-urgi.versailles.inra.fr/Seq-Repository/Annotations. Nearby genes in the linkage regions of significant SNP-trait associations with putative functions related to plant disease resistance and defense mechanisms were selected as candidates.

## RESULTS

### Seedling stage phenotypic response

In the seedling test of the landraces against of *PstS2* pathotype, 22.67% of accessions were resistant and 71.17% were susceptible and 6.17% of accessions with missing data. Out of 357 accessions of the Syrian origin, 283 accessions were susceptible. In total 110 (70%) and 27 (57%) landraces collected from Turkey and Iran, respectively, showed susceptible infection types. All the accessions from Jordan and Palestine were found susceptible while the accession from Spain and the accession from Greece showed a resistant reaction. In the case of *Warrior* pathotype, a similar trend was observed with 80% of all the accessions showing a varying degree of susceptible response and only 16% showing resistant reaction types while 4% accounted for missing genotypes. From the tested accessions against *Warrior* pathotypes, 80% of accession from Iran and Syria showed susceptible responses while 73% of the landraces collected from Turkey showed susceptible reactions. Accessions from Jordan and Palestine showed susceptible responses while the landraces from Greece and Spain showed mixed resistant and susceptible reaction types. The frequency of resistant and susceptible genotypes according to their country of origin are presented in **Fig. 1**. Minimum, maximum and mean scores for seedling and adult plant data are given in **Table 1**.

**Fig 1:**
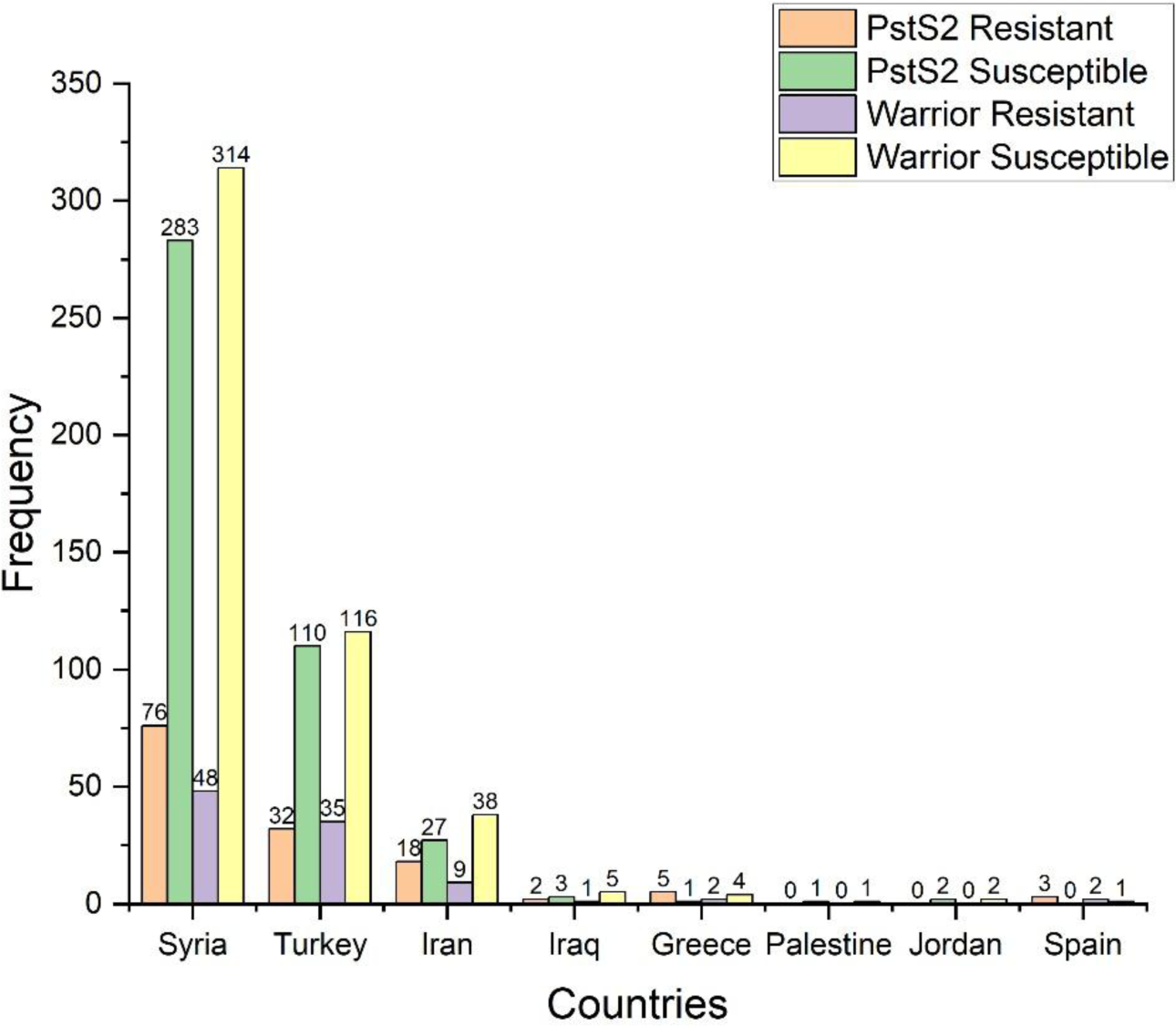
Seedling response of the 600 bread wheat landraces against PstS2 and Warrior pathotypes

### Field assessment of resistance

In adult-plant assessment, the estimates of genetic variance identified significant differences between the landraces (**Table 1**). Although variation was observed for the field responses of the tested accession during the three years, the overall field infection type patterns of the 600 landraces were consistent over the years. Highly susceptible responses of susceptible spreader indicated uniform disease spread throughout the field. During the 2016 trial, 296 (49.3%) of accessions showed resistance response (CI=0 to 20); 106 (17.6%) exhibited moderate resistance (CI=20 to 40); 130 (21.6%) showed moderately susceptible response (CI=40 to 60); and the remaining 64 (10.6%) accessions were susceptible (CI=60 to 100). During the year 2018, 53% of the genotypes showed resistance response with CI ranging from 0 to 20. 75 (12.5%) genotypes showed moderately resistant; 44 (7.3%) exhibited moderately susceptible response, and 135 (22.5%) of accessions were highly susceptible. In the year 2019, 357 (59.5%) of the genotypes showed resistance response whereas, 85 (14%) of the genotypes were highly susceptible.

Over the three years, 189 (31.5%) accessions which showed resistant response (CI= 0 to 20) in which 11 accessions showed near immune reaction type with CI less than 2; seven were found moderately resistant (CI= 20-40); two genotypes showed moderately susceptible types (CI= 40-60) while, 19 accessions were susceptible (CI= 60-100) in all the years. The year-wise frequency of disease response of the genotypes according to their geographical origins is presented in **Fig. 2**. The Range (minimum-maximum) and mean scores are given in **Table 1**.

**Fig 2:**
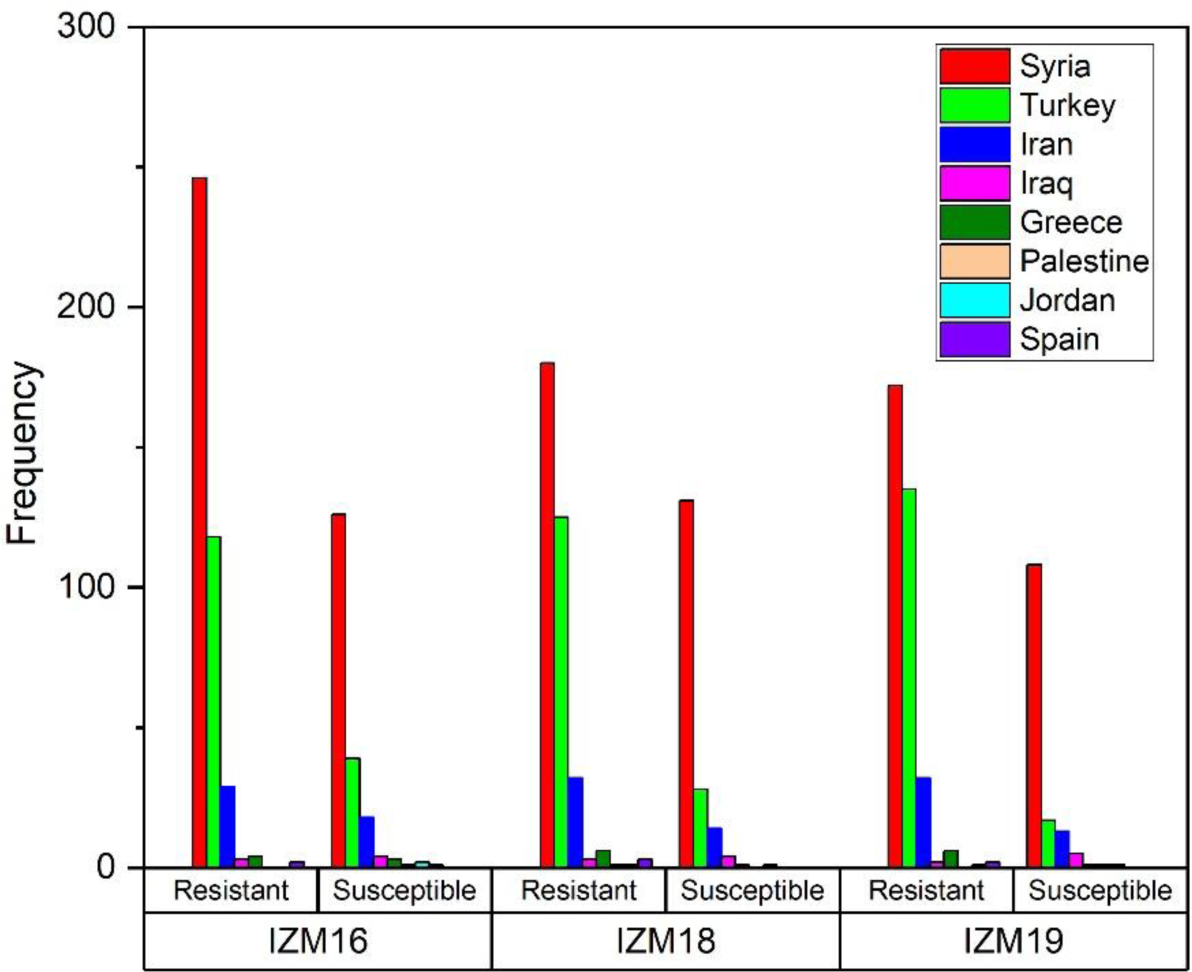
Field response of 600 bread wheat landraces over three years

### Analysis of SNP markers and LD

A total of 152K SNPs were discovered de novo. After eliminating the SNP markers call rate of less than 0.8 and minor allele frequencies of less than 0.05 (MAF <0.05) and maximum missing counts of 20%, a set of 25,169 high-quality SNP markers were used in the association analysis for resistance against the two pathotypes. The genetic framework of the polymorphic SNPs was constructed using BLAST alignment of each allele sequence with a reference genome of the Chinese Spring IWGSC RefSeq v1.0 assembly (Appels et al., 2018), resulting in 21,789 markers spread across 21 chromosomes with an average of 1000 markers per chromosome. The maximum marker density was observed on chromosome 2B with 1,477 SNPs and chromosome 4D showed a minimum markers density with 487 SNPs. The marker density for the A and B genome was almost similar with an average of 1,145 and 1,255 markers per chromosomes, respectively. However, the D genome has relatively poor marker density with an average chromosome coverage of 711 markers. Of the 25,169 markers, 3380 (13%) of the markers were not assigned to any chromosomal locations. The distribution of SNP markers across the genome is shown in **figure 3**. The extent of LD was estimated for the diversity panel using TASSEL software (Bradbury et al., 2007). It indicated the B sub-genome to have the highest LD, followed by A and D genomes, respectively. LD decreased with the increase in the physical distance between the marker loci. Average LD decay was observed after 0.4 Mb in A genome, after 0.5 Mb in B genome, and after 0.3 Mb in D genome (**Supplemental Figure S1.**).

**Fig 3:**
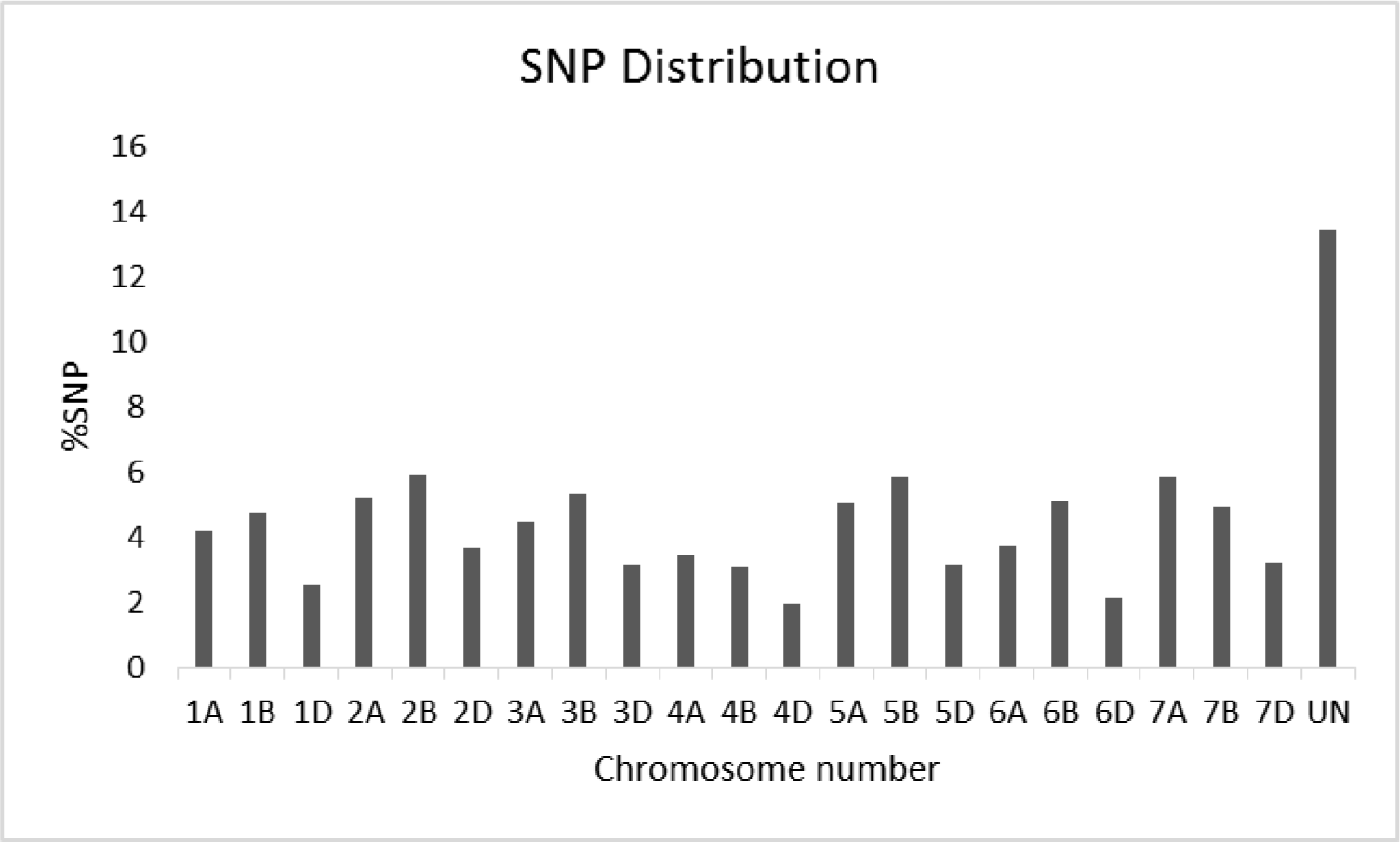
Distribution of SNPs on the chromosomes

### Population Structure Analysis

Out of the 600 bread wheat landraces, 88% (533) were collected from Syria and Turkey alone while 7% (47) were derived from Iran and the rest 3% (20) are from Iraq, Spain, Greece, Jordan and Palestine. The population structure was assessed by “Ad hoc” statistics to estimate the number of subgroups (K) based on the rate of change in the log probability of data between successive K-values. The results indicated that the population structure was best represented at K=2. In the plot of K against ΔK, there was a slope after K=2 following the flattening of the curve. This indicated that the landraces could be divided into two major groups mostly from Syria and Turkey. The results were confirmed by principal component analysis (PCA) which showed two mild population structures with admixture. The scree plot from PCA identified a steep curve at k=2 when plotted against k=1 to 30. Therefore, the population was considered to be divided into two major clusters with admixture (**figure 4**).

**Fig 4:**
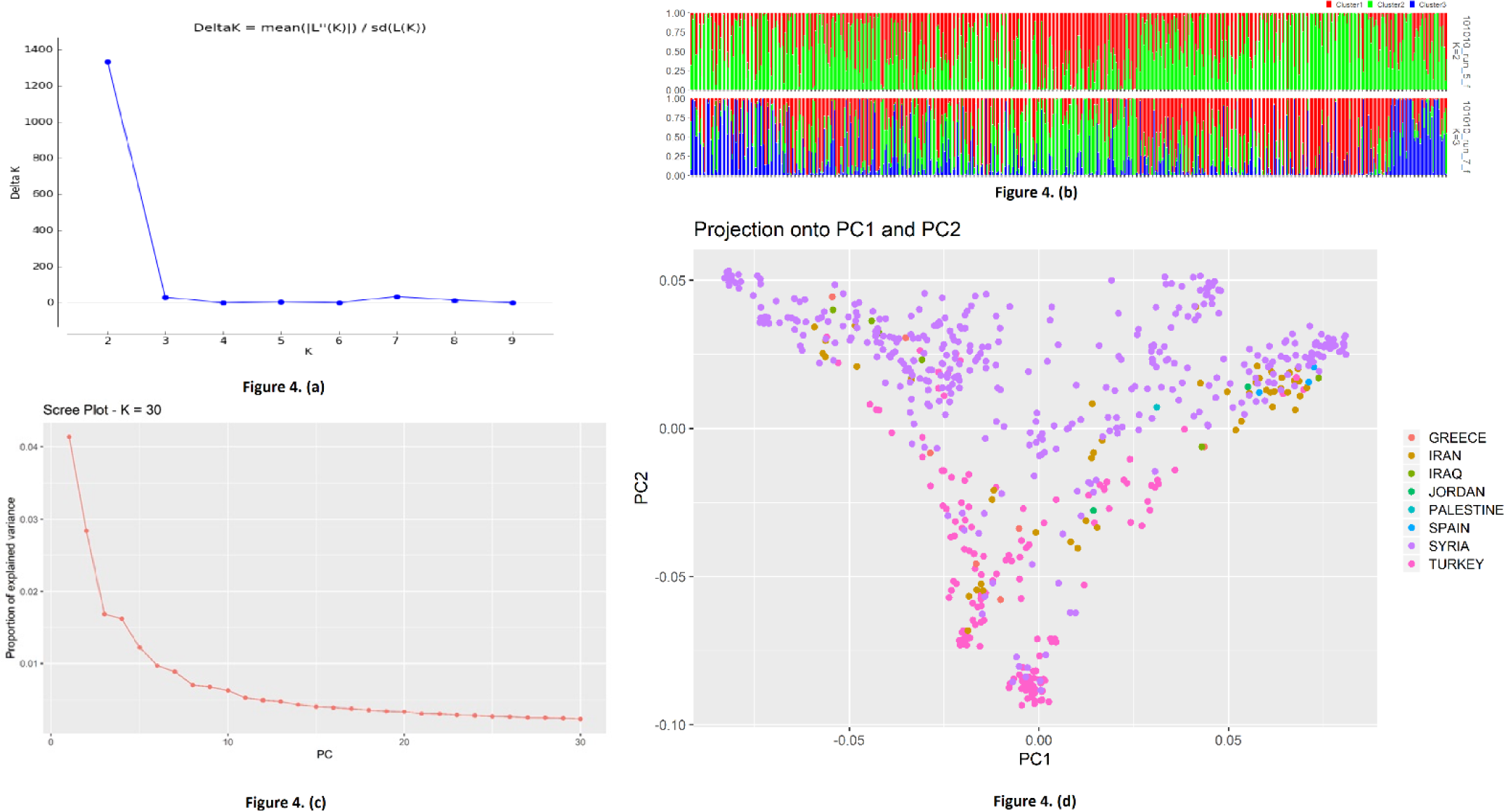
4(a). The plot of the scaled average logarithm of the probability of data likelihood [LnP(D)] and Delta K (ΔK) with K allowed to range from 2 to 10. **4(b)**. Population structure results at K=2 and K=3. Every single line represents an individual land race. **4(c)**. Scree plot derived from PCAdapt with K values ranging from 1 to 30. **4(d)**. First two Principle Components of PCA showing different population clusters with respect to the origin.

### Association Mapping Analysis

Association mapping analysis indicated potential marker-trait associations in a panel of 600 bread wheat landraces for assessment of resistance to *PstS2* and *Warrior* pathotypes of stripe rust at seedling stage and adult plant stages. Out of the 25,169 high-quality SNP markers, 21,789 (86.57%) were of known positions on the reference genome (IWGSC 2018) in which 8020, 8787 and 4982 were specific to A, B and D genomes, respectively. A total of 47 SNP markers were detected to be significantly associated with resistance to stripe rust at the adult plant and seedling stage at 19 genomic regions on chromosomes (FDR ≤ 0.05). The highest number of SNP markers were identified on chromosomes 3D and 7A i.e. 5, followed by 2D, 4B, and 5B with 4 SNPs on each of the chromosomes. No significant SNP marker was detected on chromosomes 4D and 7B. A genome had the highest number of significant SNPs i.e. 18, followed by B genome with 17 SNPs and D genome with 13 markers. For seedling stage resistance, 8 SNPs at seven genomic regions (2A, 3D, 4B, 6A, 6B, 7A and 7D) were identified to be associated with seedling stage resistance against the *PstS2* pathotype. Seedling stage analysis for *Warrior* pathotype indicated 15 significant SNP markers located on 12 genomic regions (1B, 2A, 2D, 3A, 3B, 3D, 4B, 5B, 5D, 6B, 6D and 7A) found to be associated to resistance at the seedling stage (**Table 2**). Multi-trait SNPs were detected on chromosomes 2A, 3D, 4B and 7A for adult plant resistance and seedling resistance against both pathotypes. Of the 47 significant SNP markers, 24 SNPs at 14 genomic regions were in association with resistance to stripe rust at the adult plant stage (**Table 3**). In addition, several SNPs were found significant to both adult plant and seedling stage resistances with unknown chromosomal positions, therefore, could not be co-localized with any of the previously identified *Yr* genes/QTL (**Supplemental Table S5**). Four SNPs on three genomic regions were mapped far from any previously identified *Yr* gene/QTL. Hence, these three genomic regions most likely tag new *Pst* resistance loci (**figure 5**). The remaining genomic regions putative to field and seedling resistance were mapped close to known *Yr* genes and QTL.

**Fig 5:**
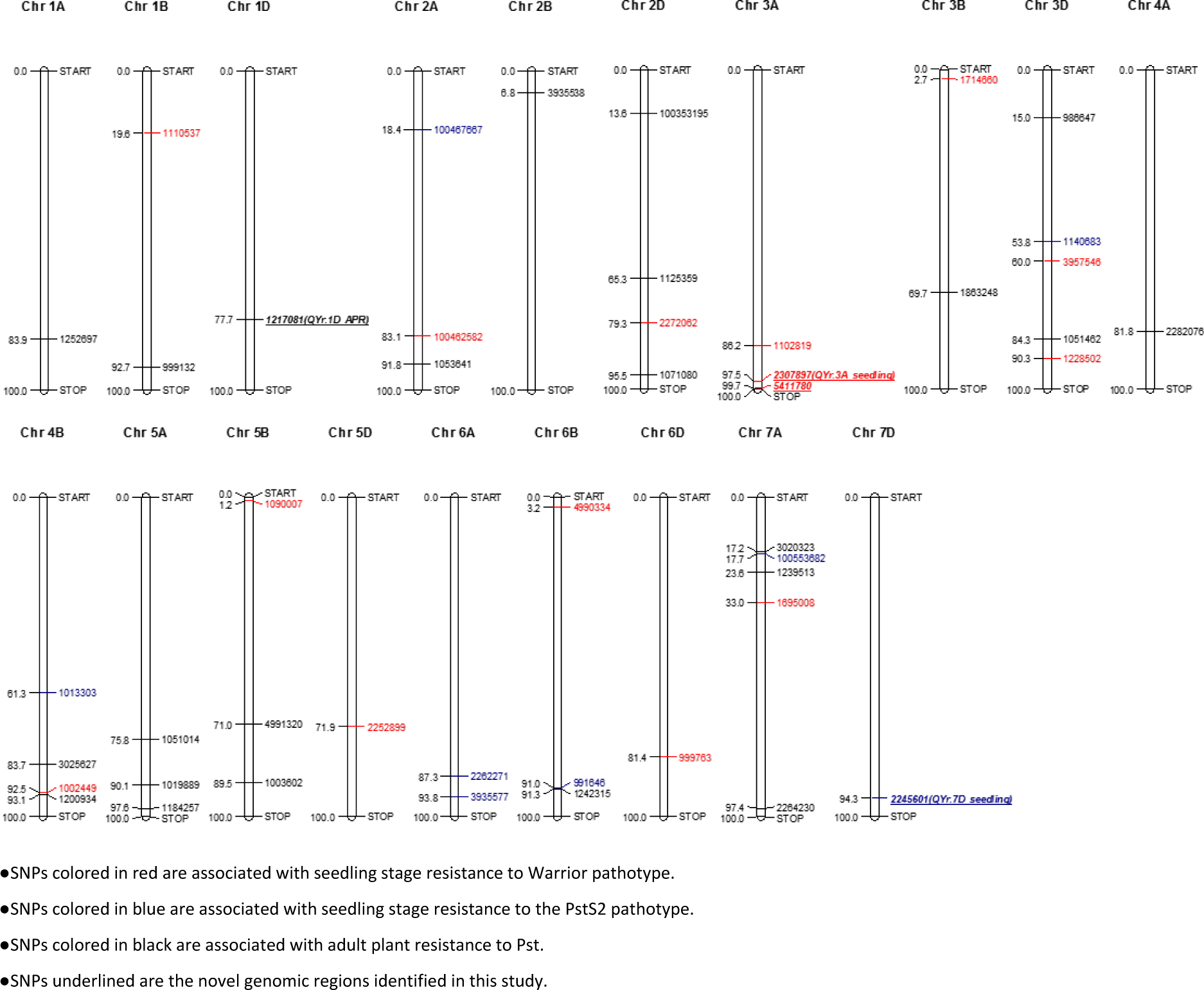
Significant SNPs detected in the study based on wheat consensus genetic map (Bulli et al., 2016)

**Table 3:**
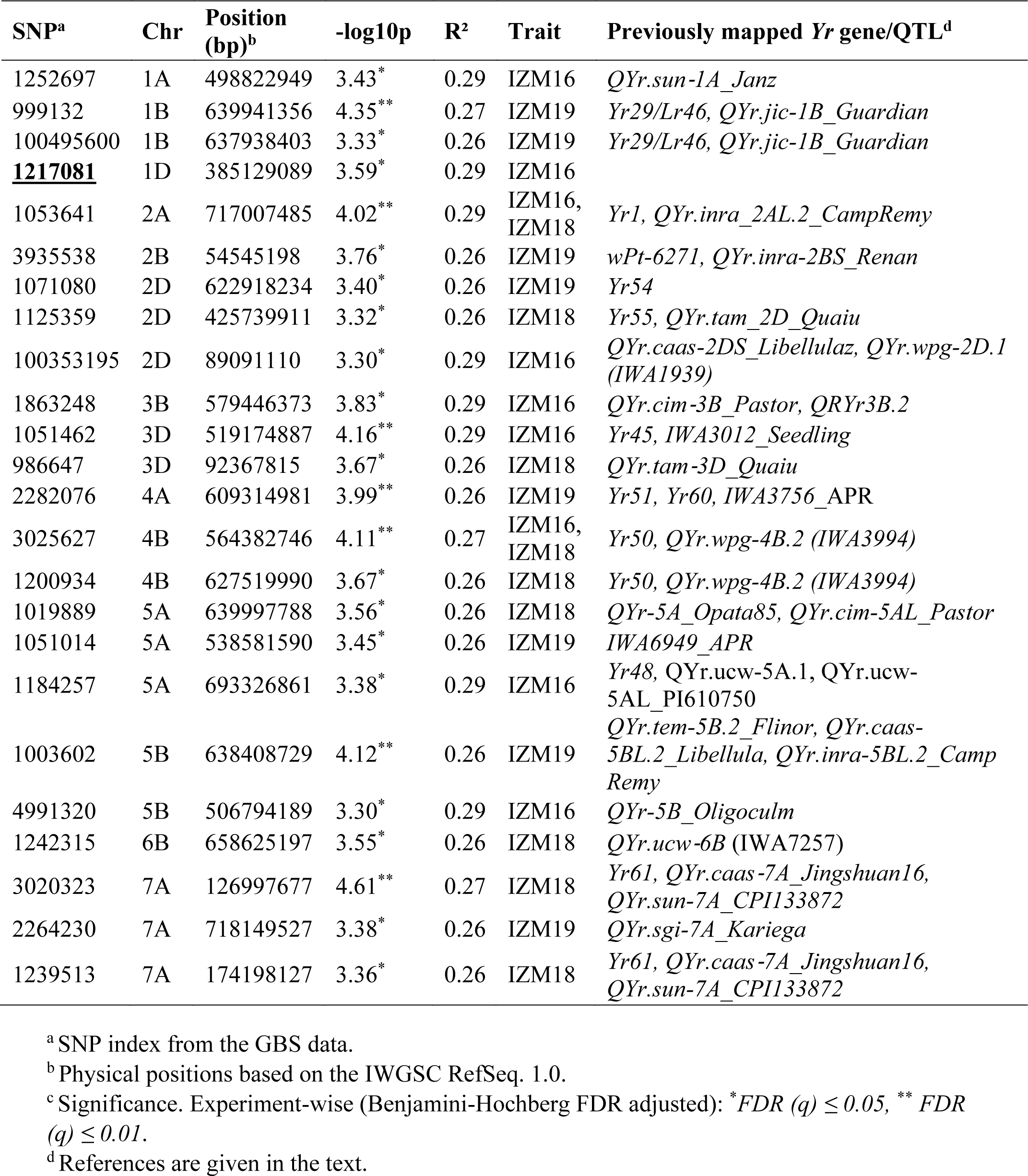
Genomic regions significantly associated with field-based Adult Plant Resistance (APR) to stripe rust in 600 bread wheat landraces collection from ICARDA gene bank

### Candidate gene predictions and function annotations

Putative genes related to disease resistance in 13 genomic regions of 24 marker-trait significant SNP loci were selected as candidates that resulted in 32 genes on the wheat reference genome assembly IWGSC *RefSeq v1.0* **(Supplemental Table S4)**. Among these, 25 genes were annotated for functional proteins involved in plant disease resistance and defense mechanism. The putative proteins/enzymes related to these genes include Leucine-rich repeat (LRR) domains, Protein kinase domain, F-box domain Proteins, Protein kinase domain coupled with Leucine-rich repeat domain (RLK), DnaJ domain proteins, Zinc finger C2H2-type proteins domain, Phosphatidylinositol-4-phosphate 5-kinases (PIP5K), Autophagy-related (ATG) protein 27, C2 domain-containing protein / VASt domain / GRAM domain-containing protein, Cytochrome P450, AP2/ERF domain Proteins, Diacylglycerol kinase-plant, ATPases Associated with diverse cellular Activities (AAA ATPases), Dehydrin-LEA (late embryogenesis abundant) proteins, BTB/POZ and MATH domain-containing protein 2 (BPM2), Xyloglucan fucosyltransferase, Legume lectin domain proteins.

## DISCUSSION

There is increased use of wheat landraces for the enhancement of genetic diversity and the mining of desirable genes (Yao et al., 2019). Therefore, wheat landraces are considered as a key genetic resource for wheat breeding (Sehgal D et al., 2016). Stripe rust caused by *Puccinia striiformis* f. sp. *tritici* is a major threat to wheat yields worldwide (Sharma Poyudal et al., 2013). Two of the most widespread pathotypes in Asia and Europe are *PstS2* and *Warrior* (D. Sharma Poyudal et al., 2013; Hovmøller et al., 2016).

### Phenotypic variability and Population structure

The AM panel consisted of 600 landraces of which 580 (96%) belonged to Syria, Turkey, and Iran alone. A total of 57 (9.5%) landraces were resistant to both the *PstS2* and *Warrior* pathotypes of stripe rust at the seedling stage out of which almost half of them i.e. 26 (45%) belonged to Syria and 19 (33%) were landraces from Turkey. The association mapping panel showed 49.3%, 53% and 59% adult plant resistance in three years respectively which is similar to the stripe rust response in Izmir to Turkish landraces reported by Sehgal et al., (2016). The population structure of 600 bread wheat landraces clustered the population into two major clusters mainly from Syrian and Turkish origin with a degree of admixture. The population’s genetic clusters found in our study are similar to the previous GWAS studies (Chen et al., 2019; Ozkuru et al., 2019; Bulli et al., 2016; Kertho et al., 2015). However, the population structure identified in this study was lesser than many previously reported GWAS studies (Liu et al., 2017; Jighly et al., 2015; Zegeye et al., 2014). This may be due to the geographic proximity of the landraces collected for this study. The regions of Turkey and Syria are part of fertile crescent which is considered as the center of origin and diversity of wheat where it has been cultivated since time unknown (Bird, 1999) therefore, the local wheat landraces evolved through time by being both adaptive to the local ecology as well as resistance to biotic and abiotic stresses. (Akcura et al., 2017). The advantage of these landraces over commercial cultivars is their cultivation over years by local farmers without interference of any scientific breeding methods lead them to adapt changing climatic conditions and evolving disease resistance and therefore it could be expected that the resistance found in these landraces is ought to be more durable and novel (Sehgal et al., 2016). Since the co-evolution of rust pathogens and wheat crop in fertile crescent for thousands of years might be the reason of durable and novel genomic resistance found in the landraces as it has been evident for durability of the transfer of common bunt resistance originated from Turkish landraces to modern cultivars saving millions of dollars for wheat industry (Bonman et al., 2006). Approximately 30% of all the landraces from Syria and Turkey showed adult plant resistance in all three years revealing high genetic diversity in both the clusters in the population. In this study, nine landraces from Turkey were found to be resistant at both seedling and adult plant stage against both *PstS2* and *Warrior* pathotypes of the stripe rust, revealing high genetic variation and usefulness of Turkish wheat landraces for stripe rust resistance. Turkish national wheat breeding program can effectively utilize landraces for biotic, abiotic stresses and quality improvement (Morgounov et al., 2016; Karagoz, 2014; Akcura, 2011).

### Significant associations in the GWAS

GWAS analysis of stripe rust using high-quality SNP markers data and *Pst* assessments from field and greenhouse experiments provided valuable information about the wheat landraces preserved in ICARDA gene bank. A total of 47 SNP markers at 19 genomic regions were detected significantly associated with experiment-wise FDR (q) ≤ 0.05 in both field and seedling experiments. Twenty-four significant SNPs identified in field experiments on 14 genomic regions on average explained 27% of the total phenotypic variation.

### Alignment of QTLs to the previously identified YR genes/QTLs

The integrated map constructed by Bulli et al., (2016) was used to compare the significant SNPs detected in the study with previously published *Yr* genes and QTL. Three genomic regions that were identified to be significant at experiment-wise FDR ≤ 0.05, SNP1217081, SNP2307897 and SNP2245601 on chromosomes 1D, 3A and 7D, respectively, were found in regions where previously no *Yr* gene/QTL was mapped. Therefore, likely tagging a new stripe rust resistance loci. A total of 47 SNPs were detected significant (FDR (q) ≤ 0.05) in seedling and field experiments, however, only the relationship of the 13 high confidence (experiment-wise FDR (q) ≤ 0.01) SNPs on 10 genomic regions with previously mapped *Yr* gene/QTL are discussed below and the remaining are presented in **Supplemental File S1**.

**Chromosome 1B.** SNP999132 and SNP100495600 detected in this study are in the same genomic regions of previously identified *Yr29* gene and a QTL (William et al., 2003; Melichar et al., 2008). However, a cultivar Lal Bahadur/Pavon 1BL harboring *Yr29* which was used in the field experiments showed resistant to moderate susceptible reaction response. *Yr29* harbors partial resistance showing slow rusting phenotype (Singh et al., 2001). Several studies have shown different phenotypic responses for *Yr29* (Willial et al., 2003; Rosewarne et al., 2006; Lillemo et al., 2008). It appears that SNP999132 is likely associated with *Yr29*.

**Chromosome 2A.** SNP1053641 and SNP4008260 (**Supplemental Table S5**) were identified on chromosome 2A which contains several *Yr* resistant genes, both SNPs seems to be in the same region as *QYr.inra_2AL.2_CampRemy* (Boukhatem et al., 2002) on the long arm of chromosome 2A. In the light of the virulence/Avirulence formula of both races it is observed that chromosome 2A is strongly associated with resistance to the *PstS2* race as *Warrior* pathotype is virulent to *Yr1*, *Yr17* and *Yr32*. Based on the consensus genetic maps of approximate positions (Bulli et al., 2016), SNP1053641 and SNP4008260 (**Supplemental Table S5**) seem to be related to the *Yr1* gene which was effective against the *PstS2* race.

**Chromosome 3D.** McIntosh et al., (2014) reported *Yr66* to be flanking at approximately 3.0cM distance to markers IWB47165 and IWB18087/IWB56281 on chromosome 3D, these markers are located at 32,220,594 bp and 3,549,510 bp/3,240,899 bp respectively. The SNP1051462 found in this study was associated with APR located at 519,174,887 bp indicating a different chromosomal region than *Yr66*. An APR gene *Yr49* and a race-specific *Yr45* gene were also mapped on chromosome 3D (McIntosh et al., 2014; Li et al., 2011). Zegeye et al., (2014) reported a stripe rust seedling resistance QTL *IWA3012_Seedling* mapped near the genomic region of SNP1051462. An SNP in association (FDR ≤ 0.05) with *Warrior* pathotype is also detected in the same region hinting the presence of a potential race-specific all stage resistance QTL. Therefore, it is likely that SNP1051462 is associated with a race-specific *Yr45* gene.

**Chromosome 4A.** Two *Yr* genes, *Yr51* and *Yr60*, have previously been reported near the genomic region of SNP2282076 associated with field APR on chromosome 4A. Randhawa et al., (2014) reported *Yr51* on chromosome 4AL as a seedling resistance gene while Herrera-Foessel et al., (2015) reported an APR gene *Yr60*. Since the region was not detected in any of the seedling experiments it is likely that SNP2282076 is related to *Yr60*.

**Chromosome 4B.** Three SNPs, SNP3025627, SNP985169 (**Supplemental Table S5)** and SNP1200934 (FDR ≤ 0.05) were found associated with APR to *Pst* on chromosome 4B. SNP985169 identified on chromosome 4B is in LD with SNP1002449 identified in the seedling experiment with *Warrior* pathotype therefore, likely tagging the same genomic region. *Yr50* and *Yr62* (Liu et al., 2013; Lu et al., 2014) and an APR QTL, *QYr.wpg-4B.2 (IWA3994)* (Naruoka et al., 2015) are mapped on this region. *Yr50* is reported as a seedling resistance gene whereas *Yr62* is reported as a High-temperature adult plant (HTAP) resistance gene which is not expressed at the seedling stage (Liu et al., 2013; Lu et al., 2014). However, this genomic region is tagged in both seedling and field experiments therefore, further analysis is required to establish its relationship with either of the earlier reported *Yr* genes. Naruoka et al., (2015) reported *QYr.wpg-4B.2 (IWA3994)* as an APR QTL therefore it is possible that SNPs identified in this study might be tagging *QYr.wpg-4B.2 (IWA3994)*. Since the region is identified in both seedling experiment with *Warrior* pathotype and field experiment it is likely that it confers an all stage race-specific QTL.

**Chromosome 5B.** Two APR and one seedling QTL were identified on chromosome 5B. SNP1003602 was found in the region of previously identified QTL *QYr.tem-5B.2_Flinor, QYr.caas-5BL.2_Libellula* and *QYr.inra-5BL.2_Camp Remy* (Feng et al., 2011; Lu et al., 2009; Mallard et al., 2005). Bansal et al., (2011) reported a genetic association between *Yr47* and *Lr52* with markers *gwm*234 and *cfb*309 in chromosome 5B which seems to be in the region of our SNP1090007. The *Yr47* is reported to confer seedling resistance therefore, it is likely that SNP1090007 is linked to *Yr47* but further analyses are required to confirm the statement. The absence of this genomic tag in seedling experiments with *PstS2* reveals the specificity of the region to *Warrior* pathotype.

**Chromosome 5D.** Stripe rust resistance gene *Yr40/Lr57* is reported on chromosome region 5D (McIntosh et al., 2014). Since there is no report of *Yr40* being susceptible or resistant to *PstS2* and/or *Warrior* pathotype and we lack the genomic information to compare our findings with the previous reports, however, based on consensus map (Bulli et al., 2016) SNP2252899 lies significantly outside the genomic region of *Yr40*. Two closest reported QTL to SNP2252899 are *QYrdr.wgp-5DL (IWA8331)* and *QYr.caas-5DL (IWA4087)_Zhong 892* (Hou et al., 2015; Liu et al., 2015). *QYr.caas-5DL (IWA4087)_Zhong 892* was reported as an APR QTL while *QYrdr.wgp-5DL (IWA8331)* as seedling resistant QTL. Since SNP2252899 is a seedling QTL furthermore, the virulence/avirulence formula of the pathotypes used was similar to *Warrior* pathotype therefore, it could be related to *QYrdr.wgp-5DL (IWA8331)*.

**Chromosome 6A.** SNP3935577 associated with the *PstS2* seedling resistant was found proximal to the genomic region of *QYr.cim-6AL_Francolin* (Lan et al., 2014). However, *QYr.cim-6AL_Francolin* is linked to a minor effect APR QTL whereas, SNP3935577 tags major effect seedling resistance to *PstS2.* Hence, it is likely that SNP3935577 and *QYr.cim-6AL_Francolin* are different. Furthermore, an adult plant resistance gene *YrLM168* (Feng et al., 2014) is at approximately 12 cM distance from SNP3935577. Additional genetic analysis is needed to confirm the relationship between SNP3935577 and *YrLM168*.

**Chromosome 6D.** The most significant SNP, SNP999763 identified to be associated with *Warrior* pathotype at the seedling stage was found on chromosome 6D near the genomic region of previously reported *QYr.ufs-6D_Cappelle-Desprez* (Agenbag et al., 2012). Agenbag et al., (2012) reported *QYr.ufs-6D_Cappelle-Desprez* as a minor effect APR QTL whereas, SNP999763 exhibit major effect seedling resistance. Zegeye et al., (2014) also reported a seedling resistance QTL IWA4455_Seedling therefore, it is likely that resistance conferred by SNP999763 is due to IWA4455_Seedling.

**Chromosome 7A.** Two Stripe rust SNPs, SNP100553682 and SNP3020323, conferring both seedling and adult plant resistance were detected in this genomic region. Two loci, SNP1695008 and SNP 1239513 significantly associated (FDR ≤ 0.05) to *Warrior* pathotype and APR were also detected in the same region. All these SNPs were detected in the genomic region of *Yr61, QYr.caas-7A_Jingshuan16* and *QYr.sun-7A_CPI133872* (McIntosh et al., 2014; Ren et al., 2012; Zwart et al., 2010). Another high-temperature seedling resistance gene *Yrxy1* (Zhou et al., 2011) was previously reported on chromosome 7A but it is unlikely that *Yrxy1* can be the source of resistance as the seedling experiments were not subjected to any high-temperature stress. The *Yr61* is reported as an all stage resistant gene (Zhou et al., 2014) and the presence of both seedling and adult plant resistance QTL in the genomic region reveals that the identified QTL in this study are related to *Yr61*.

### Novel genomic regions identified in this study and their significance

Three genomic regions were tagged where previously no *Yr* gene or QTL has been reported therefore, likely tagging novel genomic regions.

Bulli et al., (2016) reported *QYr.ucw-1D* between 24.0% to 25.6% approximate length on chromosome 1D whereas, the SNP1217081 detected in this study is located at approximately 77.7% length and no stripe rust QTL is reported in this region in any previous study indicating a novel region for resistance to stripe rust. SNP1217081 was designated as *QYr.1D_APR*.

Two markers, SNP 2307897 and SNP 5411780, located on chromosome 3A were found in association with stripe rust resistance to *Warrior* pathotype at the seedling stage. Gao et al., (2016) reported QTL associated with leaf rust response in the same chromosomal region whereas, Cavanagh et al., (2013) reported QTL associated with plant height, flowering time and thousand kernels weight indicating the potential of this region in chromosome 3A for multiple traits. However, so far no QTL associated with stripe rust is reported in this region of chromosome 3A therefore, indicating the region is a potential source for resistance to stripe rust as well. Both markers, SNP 2307897 and SNP 5411780 are tagging different genomic regions as they are not in LD with each other. The closest reported QTL *QYr.cim-3A_Avocet* (Rosewarne et al., 2012) is a minor effect QTL but based on the genetic map distances it is likely that SNPs identified in this study are tagging a novel genomic region. SNP 2307897 was designated as *QYr.3A_seedling*.

One marker SNP2245601, designated as *QYr.7D_seedling*, was found in association with seedling resistance against *PstS2* pathotype on chromosome 7D, two *Yr* resistance genes *Yr18/Lr34* and *Yr33* are reported on 7D however, according to consensus maps (Maccaferri et al., 2015; Bulli et al., 2016) SNP2245601 is outside the genomic region of any of these earlier reported genes, the approximate distance between SNP2245601 and *Yr33* at distal end is 42.5 Mb thus, based on the genetic distance of SNP2245601 it is highly likely that this SNP is indicating a novel QTL region.

In light of the scope of the disease the novel genomic regions identified here in this study are of significant importance. Many previous studies have highlighted the importance of landraces and accessions preserved in gene banks for their utilization in finding new sources of genes (Muletta et al., 2017; Naruoka et al., 2015). The identification of three novel genomic regions apart from several already reported significant genomic regions maybe the result of the large genetic diversity present in landraces used in this study. The co-evolution of landraces along with pathogen over time has enabled them to accumulate diverse resistance loci as a result making them important choice for selection in breeding programs. Although several *Yr* genes have already been reported previously for stripe rust and many of them have been validated in this study, the *Pst* pathogen has a capability of adapting and evolving continuously, building resistance against resistance genes and has been observed to cause epidemic in regions which were deemed unfavorable for it, so the currently available information, without taking anything away from its significance, is not sufficient and we need to keep mining new sources of resistance. Furthermore, the impact of rust on the agronomic traits such as yield is still extremely important therefore, exploring new sources of resistance is still of paramount importance in rust affected zones to ensure maximum wheat productivity. *PstS2* and *Warrior* are the two most widely spread pathotypes of *Pst* which are virulent to several important *Yr* genes deployed in the affected region (Tadesse et al., 2014; Hovmoller et al., 2016). Hence, the information identified here becomes of extreme significance and should/must be further investigated through molecular biology/functional genomics approaches to validate the putative candidate genes and to study their mechanism inside plant.

### Candidate Genes

Plant genes typically annotated for resistance ‘*R’* genes such as coiled-coil domain, nucleotide-binding site, leucine-rich repeat (CC-NB-LRR), receptors like proteins coupled with extracellular LRR (RLP) and kinase domain coupled with LRR (RLK) were selected as candidate genes along with genes with putative functions related to plants defense mechanism and signaling pathways and disease resistance. The typical *R* genes harboring LRR, RLK, LRR-NB-ARC, LRR-RLK and F-box/LRR domain proteins were identified and selected as candidate genes on seven genomic regions linked to ten SNPs (**Supplemental Table S4**). LRRs occur in proteins ranging from viruses to eukaryotes and are involved in disease resistance, apoptosis, and immune response and are typically annotated for *R* genes (Enkhbayar et al., 2004; Kobe & Kajava 2001; Rothberg et al., 1990). Amongst these ten SNPs in linkage with typical *R* genes, SNP5411780 is a novel SNP detected in this study on chromosome 3A and is linked with resistance to *Warrior* pathotype at the seedling stage. Another gene *TraesCS3A01G537200* is associated with SNP5411780, *TraesCS3A01G537200* encodes Dirigent Protein-Plant disease resistance response proteins which are induced during disease response in plants (Seneviratne et al., 2015; Pickel et al., 2010). The presence of the *R* gene associated with the novel QTL detected in this study on chromosome 3A validates the presence of potential QTL/*Yr gene(s)* in the region. Two putative genes *TraesCS1D01G286800* and *TraesCS3A01G512600* with Protein kinase-like domain were found in association with SNP1217081 and SNP2307897 on chromosome 1D and 3A. Protein kinase-like domains are involved in apoptosis, signaling and regulatory pathways and are typically annotated for disease resistance genes (Engh & Bossemeyer 2002; Walker et al., 2000). The APR QTL, SNP1217081 tags a novel genomic region thus, the presence of a candidate gene with putative functions involving disease resistance and defense mechanisms further confirms the presence of a novel candidate APR QTL. A novel QTL (SNP2245601) on chromosome 7D associated with seedling resistance to *PstS2* was found in linkage with two candidate genes *TraesCS7D01G494000* and *TraesCS7D01G494200* with putative stress and disease resistance related proteins respectively. *TraesCS7D01G494200* contains the Legume lectin domain which is believed to play a role in the protection against pathogens (Roopashree et al., 2006; Loris et al., 1998). The presence of these two genes highlights the importance of this novel genomic region identified in the current study. A putative candidate gene *TraesCS4B01G190400* that translates Autophagy-related (ATG) protein 27 is in association with seedling resistance to *PstS2*. The autophagy-related proteins play important role in resistance response to fungus infection, such as *Blumeria graminis* f. sp. *tritici* as well as in the elimination and suppression of invading bacteria (Pei et al., 2014; Yoshimori & Noda 2008). *TraesCS6A01G306300* encoding AP2/ERF domain was found in linkage with SNP2262271 which is associated with *PstS2* seedling resistance. AP2/ERF is found in various pathogenesis-related (PR) defense genes (Magnani et al., 2004). TraesCS6A01G347800 was identified in the region of SNP3935577 and was annotated for Diacylglycerol kinase. This enzyme regulates plant signaling pathways (Gomez-Merino et al., 2005). Putative candidate gene *TraesCS2A01G479000* encoding Phosphatidylinositol-4-phosphate 5-kinases (PIP5K) was found in association with SNP1053641. PIP5K plays an important role in coordinating plant growth in response to external factors. (Lou et al., 2007; Mikami et al., 1998). SNP3020323 was in association with *TraesCS7A01G171600* regulating the Xyloglucan fucosyltransferase enzyme. Cell walls in plants play vital roles in disease resistance, signal transduction and development in plants and fucosyltransferase adds an important fucosyl residue in the cell wall which helps cell wall keep regulating all those functions (Perrin et al., 1999). Five genes (*TraesCS1A01G306700, TraesCS3D01G447900, TraesCS3D01G239300, TraesCS3D01G354100LC, and TraesCS5D01G310700*) on three genomic regions (1A, 3D and 5A) encoded F-box domain proteins. F-box proteins are involved in plant vegetative and reproductive growth and development. These proteins are reported to regulate cell death and defense when the pathogen is recognized in the Tobacco and Tomato plant (Peng et al., 2012; van den Burg et al., 2008).

## CONCLUSION

The results of the current study emphasize the prospects of taking advantage of high genetic diversity in bread wheat landraces mainly due to historical recombination. The landraces preserved at the ICARDA gene bank possess a wide range of phenotypic diversity for both the seedling stage and APR to wheat stripe rust. Accessions with higher resistance response across seedling and adult-plant stage can be a vital genetic material in breeding programs against stripe rust and the molecular markers linked to resistant QTLs could serve as reliable breeding tools for future stripe rust breeding programs. The *in silico* analysis identified the proteins, involved in plants disease resistance and defense mechanism, linked to previously reported and novel QTL detected in this study thus, further confirming the significance of the study and the genomic regions harboring these resistance loci. This *in silico* functional analysis provided the baseline for the next phase of the study which is to utilize the marker sequences and convert them into functional markers. Which then will be utilized to validate the QTL by using bi-parental populations, Recombinant inbred lines (RILs) or near-isogenic lines (NILs).

## Supplemental Information

Supplemental Table S1. List of bread wheat landraces collected from ICARDA gene bank used in the study including their country of origin.

Supplemental Table S2. Virulence/Avirulence formula for the *PstS2* and *Warrior* Pathotypes of

*Pst*.

Supplemental Table S3. Differential set used in the study for race typing.

Supplemental Table S4. Candidate genes with putative enzymes/proteins associated with significant SNPs.

Supplemental Table S5. List of SNPs associated (marker-wise p ≤ 0.001) to both seedling and field-based resistance to stripe rust.

Supplemental Fig. S1. LD decay plots of wheat genome A, B, and D.

Supplemental Fig. S2. 600 X 600 kinship matrix based on simple matching genetic similarities (IBS, identity by state).

Supplemental File. S1. Relationship of significant genomic regions (FDR (q) ≤ 0.05) associated with seedling and field resistance with previously identified *Yr* genes or QTL.

## ACKNOWLEDGMENTS

We thank Dr. Awais Rasheed (CIMMYT) and Ms. Pinar Guner (EGE University) for their valuable advice and support. This work was supported by The Scientific and Technological Research Council of Turkey (TUBITAK) with the project number of 117O049.

## Supplementary Files Information

**Supplementary Figure S1:** LD decay plots of wheat genome A, B, and D.

**Supplementary Figure S2:** 600 X 600 kinship matrix based on simple matching genetic similarities (IBS, identity by state).

**Supplementary Table S1:** List of bread wheat landraces collected from ICARDA gene bank used in the study including their country of origin.

**Supplementary Table S2:** Virulence/Avirulence formula for the PstS2 and Warrior Pathotypes of Pst.

**Supplementary Table S3:** Differential set used in the experiments for Pst pathotype identification.

**Supplementary Table S4:** Candidate genes with putative enzymes/proteins associated with significant SNPs

**Supplementary Table S5:** List of SNPs associated (marker-wise p<0.001) to both seedling and field-based resistance to stripe rust.

**Supplementary File S1:** Relationship of significant genomic regions (FDR (q) ≤ 0.05) associated with seedling and field resistance with previously identified *Yr* genes or QTL.

